# SEGS-1 episomes generated during cassava mosaic disease influence disease severity

**DOI:** 10.1101/2024.06.12.598742

**Authors:** Evangelista Chiunga, Catherine D. Aimone, Cyprian Rajabu, Mary M. Dallas, Josep Ndunguru, José T. Ascencio-Ibáñez, Elijah M. Ateka, Linda Hanley-Bowdoin

## Abstract

Cassava is an important root crop that is produced by smallholder farmers across Sub-Saharan Africa. Cassava mosaic disease (CMD), which is caused by a complex of cassava mosaic begomoviruses (CMBs), is one of the most devastating diseases of cassava. A previous study showed that SEGS-1 (sequences enhancing geminivirus symptoms), which occurs in both the cassava genome and as an episome during CMD, can increase CMD disease severity and overcome host resistance. In this report, we examined the effects of exogenously applied SEGS-1 on the incidence of CMB infection, symptom severity, and viral DNA copy number in five cassava cultivars that ranged from highly susceptible to highly resistant to CMD. These studies revealed that the effect of SEGS-1 is cultivar dependent. Susceptible cultivars developed severe CMD in the absence or presence of exogenous SEGS-1, while exogenous SEGS-1 increased disease severity in cultivars carrying CMD2 but not CMD1 resistance. Analysis of infected plants in the absence of exogenous SEGS-1 revealed that some, but not all cultivars, form SEGS-1 episomes during CMD. The presence of endogenous SEGS-1 episomes in TME14, a CMD2 resistant cultivar, correlated with CMD severity. In contrast, TME3, a closely related CMD2 cultivar, did not produce endogenous SEGS-1 episomes and showed more resistance than TME14. DNA sequence analysis indicated that the different capacities of TME3 and TME14 to form SEGS-1 episomes is unlikely due to sequence differences in and around their genomic SEGS-1 loci. Because of its inability to form episomes, TME3 was used to map the functional regions of SEGS- 1 to sequences flanking the epsiome junction, but junction itself was not required for activity. Together, these experiments provided insight into the functional form of SEGS-1 in cassava and the effect of cassava genotype on SEGS-1 activity.

## Importance

Cassava is the number one crop produced in Africa as source of food and income for most smallholder farmers. Globally, cassava is one of the most important starch sources and the demand for its production has increased. Its ability to withstand both drought and poor soil conditions has made cassava an excellent crop for providing sustainable food security and, hence, to reduce hunger and poverty, especially in the developing world. However, cassava is impacted by cassava mosaic disease (CMD), which can cause up to 100% yield losses. Circular single-stranded DNAs like SEGS-1 and the novel satellite SEGS-2 have been associated with increased geminivirus symptom severity. This study identified the regions of SEGS-1 responsible for breaking CMD2 resistance in cassava. It also showed that endogenous SEGS-1 episomes can be generated by a CMD2 resistant cultivar and that episomal presence correlates with disease severity. All cassava cultivars have a copy of SEGS-1 in their genomes which has the potential to negatively impact the development of stable CMD resistance in cassava.

## Introduction

Cassava (*Manihot esculenta* Crantz) is an important staple crop in many tropical and subtropical regions of the world (Bayata, 2019). More than half of cassava produced worldwide is grown in Sub-Saharan Africa region (FAOSTAT, 2021) where it is cultivated primarily by smallholder farmers. Cassava is a drought tolerant crop and can grow at high temperatures, but its production can be severely limited by viral disease losses (Okogbenin et al., 2013). Cassava mosaic disease (CMD) is endemic across the African continent and causes substantial crop losses on smallholder and commercial production farms (Legg et al., 2011). CMD is caused by 11 DNA viruses collectively known as cassava mosaic begomoviruses (CMBs) (Crespo-Bellido et al., 2021). Nine CMB species occur in Africa, where they are transmitted by whiteflies (*Bemisia tabaci* Genn.) and by propagation of infected stem cuttings (Legg et al., 2015).

*Begomoviruses* comprise the largest genus in the *Geminiviridae*, a family of plant viruses with single-stranded DNA (ssDNA) genomes that are encapsidated into double icosahedral particles (Hanley-Bowdoin et al., 2013). CMBs have bipartite genomes that consist of two small, circular DNA molecules designated as DNA-A and DNA-B, both of which are required for systemic infection (Stanley & Gay, 1983). Together, the two genome components encode 8-9 canonical proteins that are involved in viral replication, transcription, encapsidation, movement, and countering host defenses (Yang et al., 2016). The viral proteins interact with diverse host proteins to perform their functions (Hanley-Bowdoin et al., 2013). Recent studies have suggested that the coding capacity of begomoviruses is not limited to the canonical open reading frames (ORFs), and that the number of viral proteins may be greater due to the presence of previously unidentified ORFs and alternative splicing (Gong et al., 2021).

Begomovirus genomes undergo mutation (nucleotide substitution and small indels), recombination and reassortment (Aimone et al., 2021b; Crespo-Bellido et al., 2021; Duffy & Holmes, 2009). CMBs often occur in coinfections in which different viruses can act synergistically and cause more severe disease (Pita et al., 2001). It is likely that the different CMB species arose via recombination between coinfecting viruses (Crespo-Bellido et al., 2021). In the 1990s and 2000s, synergy between African cassava mosaic virus (ACMV) and a recombinant CMB contributed to a severe CMD pandemic that spread from Uganda to other sub-Saharan countries (Deng et al., 1997; Pita et al., 2001; Zhou et al., 1997). In response to the pandemic, many African farmers adopted cassava cultivars with the CMD2 locus, which confers resistance to CMBs (Akano et al., 2002; Rabbi et al., 2014). Recent studies have indicated that CMD2 resistance is due to mutations in the cassava gene encoding the catalytic subunit of DNA polymerase (Lim et al., 2022).

Two novel DNAs, designated SEGS-1 (sequences enhancing geminivirus symptoms; DNA-II; GenBank accession no. AY836366) and SEGS-2 (DNA-III; AY836367), were amplified from cassava plants showing severe CMD symptoms in Tanzania (Ndunguru et al., 2016). Laboratory studies showed that the presence of either SEGS-1 or SEGS-2 increases CMD symptom severity and that SEGS-1 can overcome CMD2 resistance (Ndunguru et al., 2016). SEGS-1 and SEGS-2 only show 23% overall sequence identity but display 99% and 84-87% identity, respectively, to sequences in the cassava genome. All cassava genomes examined to date encode a full-length copy of SEGS-1, but no full-length copy of SEGS-2 has been found. SEGS-1 and SEGS-2 also occur as small, circular DNA episomes in infected cassava plants. SEGS-2, but not SEGS-1, has been detected in virions in infected cassava and viliferous whiteflies, suggesting that SEGS-1 and SEGS-2 are functionally distinct.

Begomovirus infection in Arabidopsis is also enhanced by SEGS-1 and SEGS-2 when either is provided as exogenous DNA or as a transgene (Aimone et al., 2021a). Experiments in Arabidopsis established that SEGS-2 activity is dependent on a small open reading frame, that SEGS-2 occurs as both double and single-stranded episomes in ACMV infected plants, and that the single-stranded form of SEGS-2 is packaged into virions. These results, in combination with the results in cassava and experiments in tobacco cells showing that SEGS-2 replicates in the presence of ACMV DNA-A, establish that SEGS-2 is a novel satellite (Aimone et al., 2021a; Ndunguru et al., 2016). In contrast, there is no evidence that SEGS-1 forms single or double-stranded episomes on Arabidopsis or replicates in tobacco cells co-transfected with ACMV DNA-A. Instead, a linear copy of SEGS-1 integrated into the Arabidopsis genome enhances ACMV infection (Rajabu et al., 2023), raising the possibility that the full-copy of SEGS-1 in the cassava genome can also enhance CMB infection during CMD.

In the study reported here, we examined the effects of exogenous and endogenous SEGS-1 sequences on CMB infection in five cassava cultivars that ranged from highly susceptible to highly resistant to CMD. We asked if there is a correlation between the presence of SEGS-1 episomes and disease severity, determined the regions of SEGS-1 that mediate disease enhancement, and assessed the role of episome junction sequences on SEG-1 activity. Together, these experiments provided insight into the functional form of SEGS-1 in cassava and the effect of cassava genotype on SEGS-1 activity.

## Materials and Methods

### SEGS-1 constructs

The clones corresponding to the SEGS-1 monomer (S1-1.0; pNSB2000), and a SEGS-1 partial dimer with two copies of the GC-rich region (S1-1.5a; pNSB1829) were described previously (Rajabu et al., 2023) .SEGS-1 regions G (positions 1–277), I (positions 275–646), J (positions 644–756), and F (positions 757–1007) (Supplementary Figure 1) were amplified in reactions containing S1-1.0 plasmid DNA as template. SEGS-1 region N was generated by cleaving the *Kpn*1 site using the Q5-site directed mutagenesis protocol (New England Biolabs, Ipswich, MA) from S1-1.5a (pNSB1829) and joining SEGS-1F and SEGS-1G using the indicated primers in Supplementary Figure 1 and the Q5 high-fidelity DNA polymerase (New England Biolabs, Ipswich, MA). All the primers except for those for site-directed mutagenesis added *Not*I restriction sites at the 5’ and 3’ends of PCR products. G and F also included short sequences (G: 3 nt at 5’ end; F: 13 nt at 3’ end) that flank the cloned SEGS-1 sequence and are not present in the cassava genomic sequence. The PCR program consisted of an initial denaturation step at 98 °C for 30 seconds, followed by 30 cycles of (98°C for 10s, annealing temperature for 30s, and 72°C for 2 min) and a final extension step (72°C for 2 min). Products were resolved on a 1% (w/v) agarose gel and gel purified using a QIAquick Gel Extraction kit (Qiagen, Hilden, Germany) according to manufacturer’s instructions. Purified PCR products were digested with *Not*I (New England Biolabs) and ligated into pMON721 linearized with *Not*I to generate the SEGS-1 G, I, J, F and N clones (Supplementary Figure 1). All clones were confirmed by Sanger sequencing.

### Cassava infection

Cassava plants (*Manihot esculenta* cv. Namikonga, Kibaha, TME14, TME3, and TMS300572) were propagated from stem cuttings and grown at 28°C under a 12-h light/dark cycle. Plants with ca. 8-10 nodes and stems 1.5 cm in diameter (ca. 2 months after propagation) were inoculated at the apical meristem area using a hand-held micro sprayer (40 psi) to deliver gold particles coated with four plasmid DNAs (100 ng/plasmid/plant) (Aimone et al., 2022; Aimone et al., 2021b; Ascencio-Ibáñez et al., 2023). The clones contained partial tandem dimers corresponding to DNA- A or DNA-B of African cassava mosaic virus (ACMV; GenBank accessions MT858793.1 and MT858794.1) or East African cassava mosaic Cameroon virus (EACMCV; AF112354.1 and FJ826890.1) (Hoyer et al., 2020). ACMV and EACMCV were co-inoculated to generate coinfections in the absence or the presence of a SEGS-1 plasmid DNA.

Experiments characterizing the effect of SEGS-1 on viral infection in different cultivars included 30 plants for each infection treatment (ACMV+EACMCV or ACMV+EACMCV+S1- 1.5a) and 3 mock-inoculated plants as negative controls. Experiments characterizing the impact of different SEGS-1 fragments on TME3 infection included 10 plants for each infection treatment and 3 mock-inoculated plants. All experiments were repeated a minimum of 2 times with similar results. Disease symptoms were monitored visually at 28 days post inoculation (dpi). Symptoms were scored using a disease severity scale (scale: 1 = no symptoms to 5 = very severe) in new growth (Supplementary Figure 2). The percent symptomatic plants was calculated as the percent of plants with symptom scores ≥ 2 relative to the total number of inoculated plants within a treatment. Average symptom scores were determined using all scores ≥ 2 within a treatment. The percent symptomatic plants and average symptom scores were compared between treatments within an experiment using Chi-square and Wilcoxon ranked sum tests, respectively, and p-values < 0.05.

### DNA extraction, viral DNA quantification, and SEGS-1 episome analysis

Six leaf punches from the base of the second visible leaf (L2) relative to the top of a cassava plant were sampled at 28 dpi, flash-frozen in liquid nitrogen, and stored at −80°C. The frozen tissue was pulverized using a homogenizer (MM 301, RETSCH-Laboratory Mills, Clifton, NJ), and total DNA was extracted using the MagMax^TM^ Plant DNA Isolation Kit according to manufacturer’s instructions (Thermo Fisher Scientific, Waltham, MA). The copy numbers of ACMV DNA-A and EACMCV DNA-A were measured by quantitative PCR (qPCR) in total DNA samples (10 ng/assay) as described previously (Aimone et al., 2022).The ACMV A primer pair (ACMV-MSC1 and CMA-rev4) (Supplementary Table 1) was used to generate the qPCR results in Figure 1. The ACMV-A primer pair (P3P-AA2F and P3P-AA2R+4R; Supplementary Table 1) was used to generate the ACMV DNA-A qPCR data for Figure 2, 4, 5, and 6. EACMCV DNA-A was quantified using the primer pair – EACMVQ1 and EACMVQ2 (Supplementary Table 1). The qPCR conditions, standard curves, and calculation of DNA-A copy number/ng of total DNA were described by (Aimone et al., 2022).Viral DNA copy number was compared between treatments within an experiment using a two-tailed Students’ T test and a p-value < 0.05.

**Figure 1.**
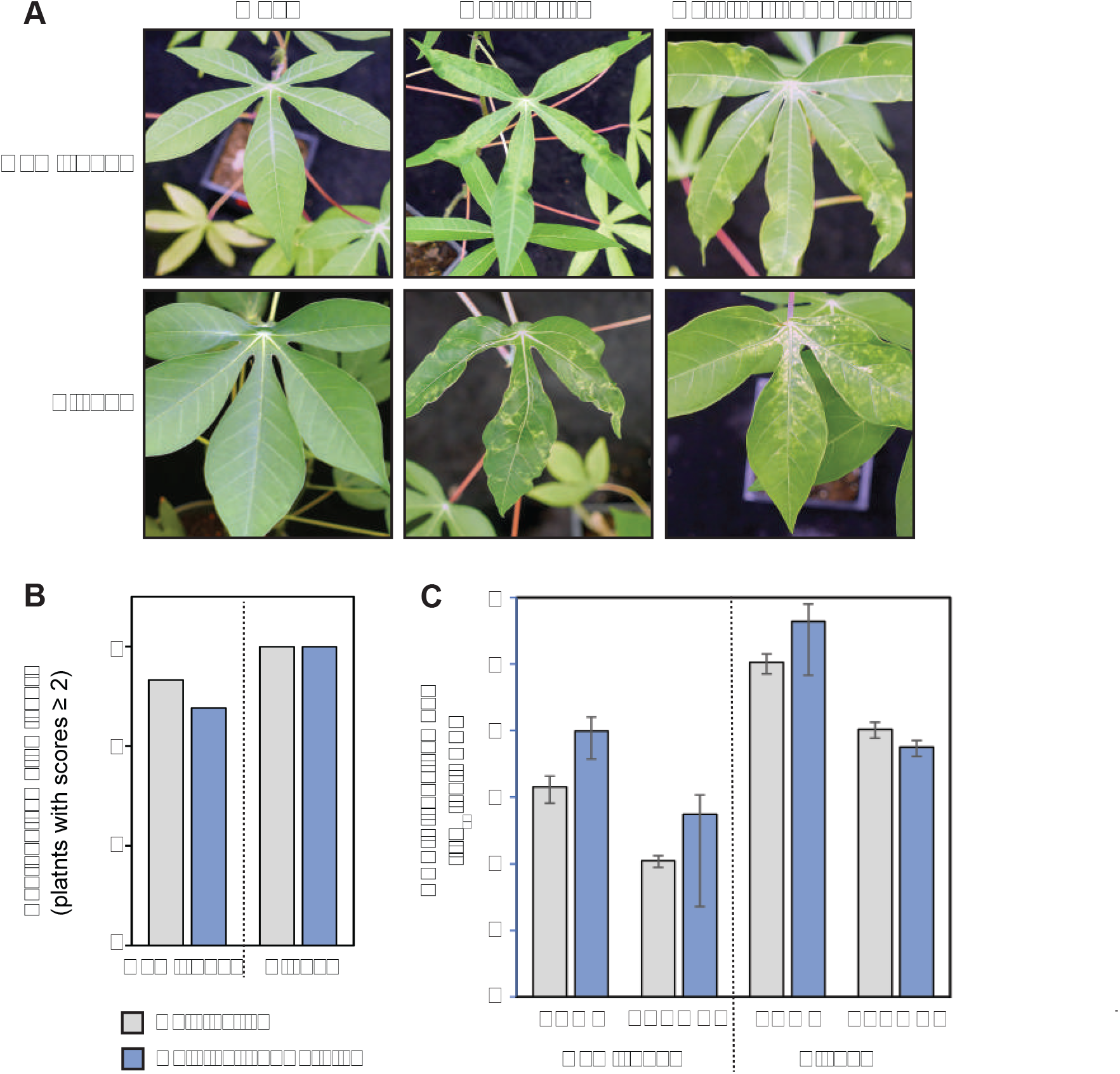
Exogenous SEGS-1 does not enhance symptoms in CMD susceptible and tolerant cassava. Cassava cultivars Namikonga (susceptible) and Kibaha, (tolerant) were co-inoculated with ACMV+EACMCV alone (coinfection) or in combination with exogenous SEGS-1 DNA (coinfection + S1-1.0) under controlled conditions and monitored for incidence of infection, symptom severity, and viral copy number at 28 dpi. **(A)** Images of leaves from the susceptible and tolerant cassava cultivars. Mock-inoculated controls are shown on the left. **(B)** Average symptom scores of plants with symptom scores ≥2. **(C)** Average copy number of viral DNA-A per ng total DNA on a log_10_ scale. The bars represent 2 standard errors. No significant differences were detected in the absence or presence of exogenous SEGS-1 DNA in **(B)** using a Mann Whitney test or in **(C)** using a two-tailed Students’ T test.

**Figure 2.**
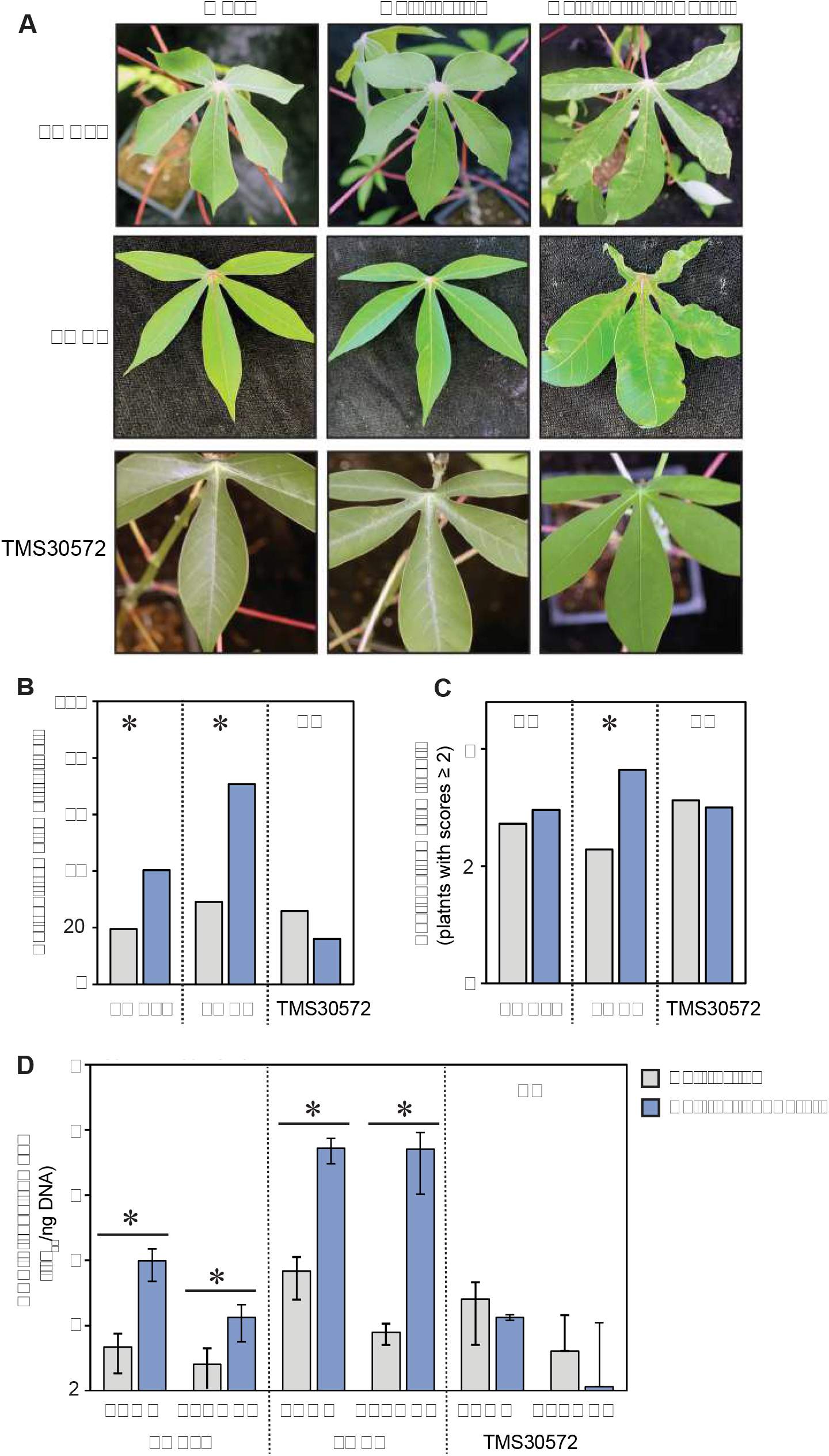
Exogenous SEGS-1 can overcome CMD2 resistance in cassava. The CMD resistant cultivars, TME-14 (CMD2), TME3 (CMD2), and TMS300572 (CMD1) were co-inoculated with ACMV+EACMCV alone (coinfection) or in combination with exogenous SEGS-1 DNA (coinfection+S1-1.0) under controlled conditions and monitored for incidence of infection, symptom severity, and viral copy number at 28 dpi. **(A)** Images of leaves from the resistant cassava cultivars. Mock-inoculated controls are shown on the left. **(B)** Incidence of plants (%) with symptom scores ≥2. **(C)** Average symptom scores of plants with symptom scores ≥2. **(D)** Average copy number of viral DNA-A per ng total DNA on a log_10_ scale. The bars represent 2 standard errors. Asterisks indicate significant differences (p-value<0.05) between coinfection and coinfection+S1.0 treatments within a cultivar. A Mann Whitney test was used in **(B)** and **(C)**, and a two-tailed Students’ T test was used in **(D)**.

**Figure 3.**
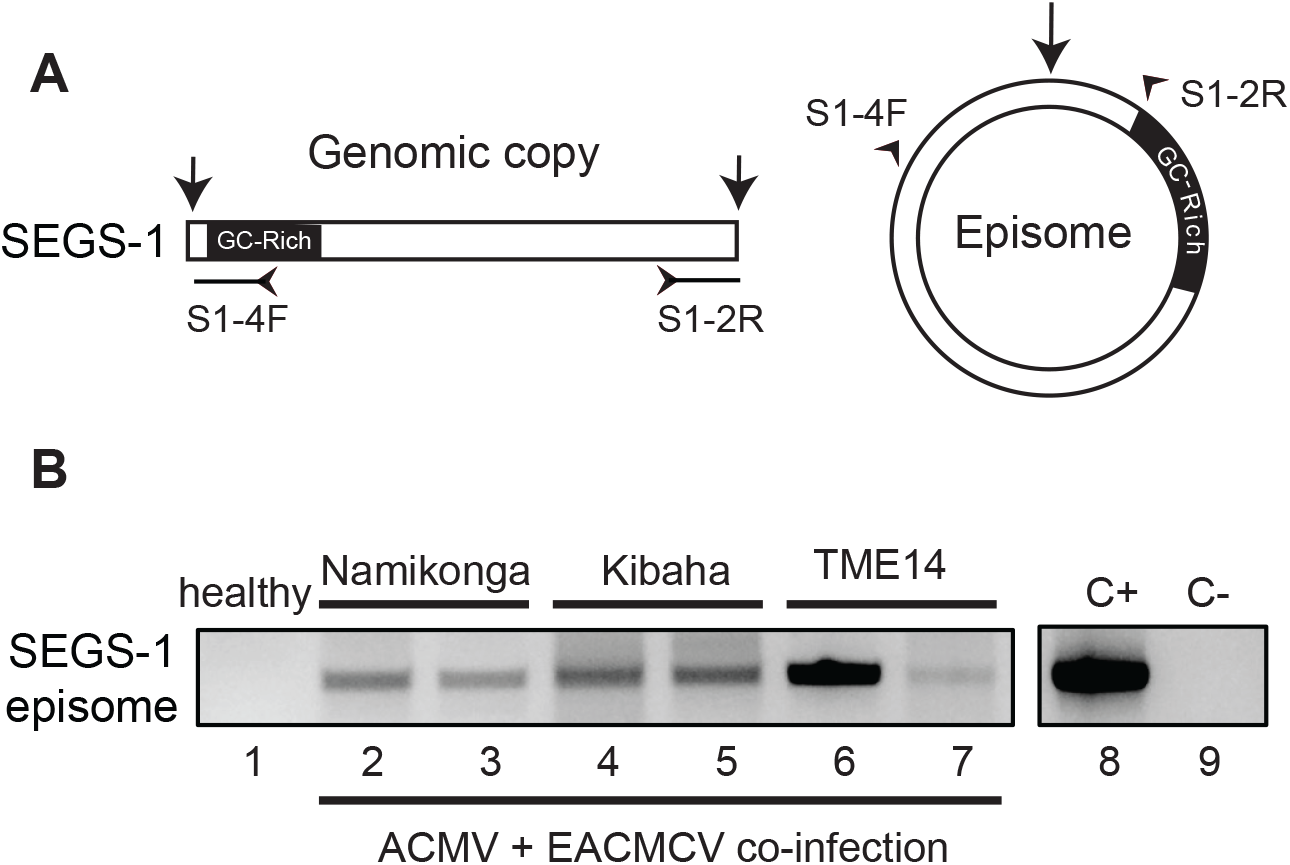
Formation of endogenous SEGS-1 episomes during CMD. **(A)** Diagram of divergent primers that amplify across the junction of a SEGS-1 episome but not the linear SEGS-1 sequence in the cassava genome. **(B)** SEGS-1 episomes were detected in Namikonga (susceptible, lanes 2 and 3), Kibaha (tolerant, lanes 4 and 5), and TME-14 (CMD2 resistant) plants co-inoculated with ACMV+EACMCV alone at 28 dpi. No episomes were detected in the healthy cassava control (lane 1), demonstrating that the divergent primers do not amplify the SEGS-1 genomic sequence. Lane 8 is the positive control (+C) using plasmid S1-1.5a, that contains the junction sequences as template. Lane 9 is the no template control (-C).

**Figure 4.**
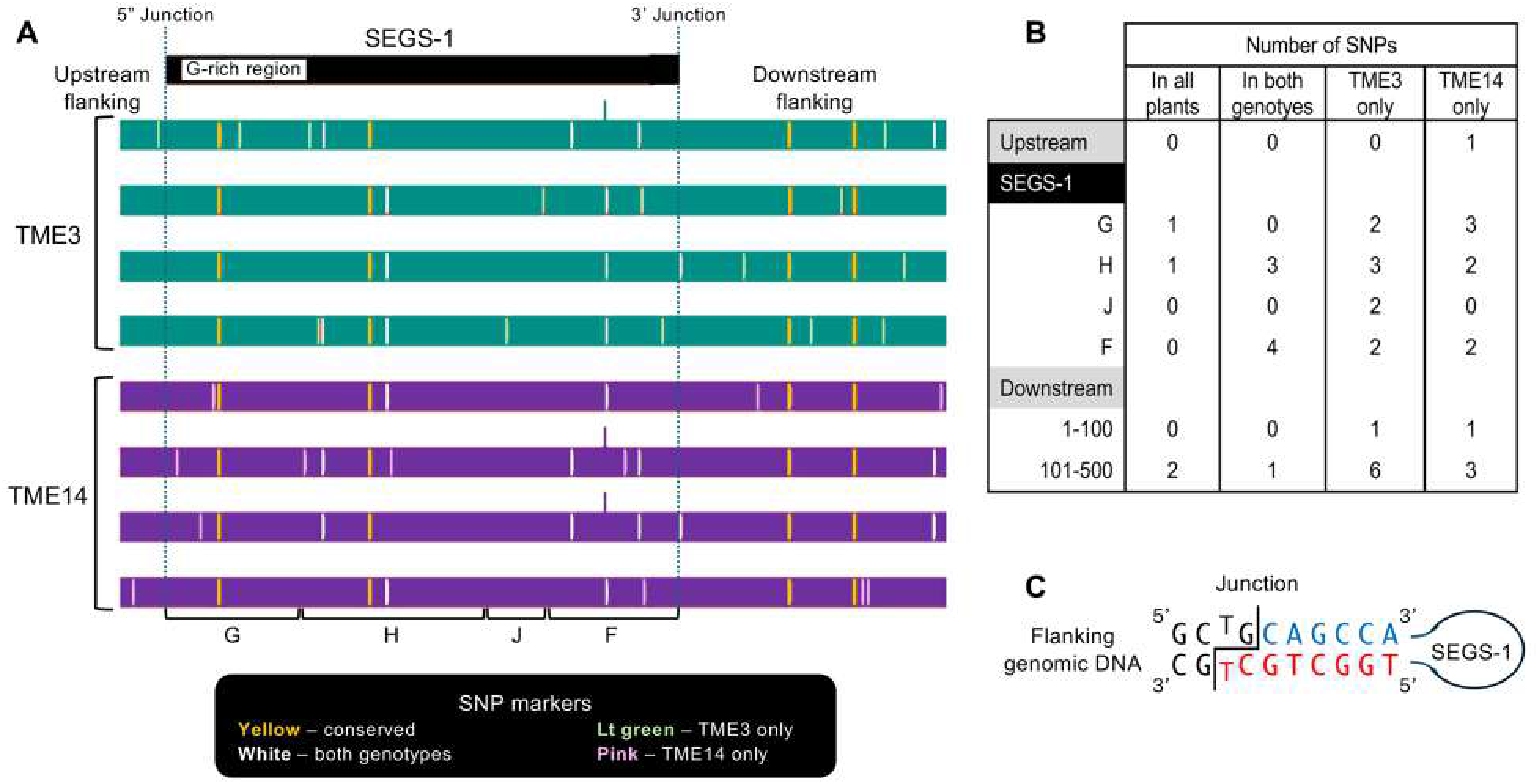
Sequence comparison of SEGS-1 genomic regions in CMD2 resistant cultivars. The genomic copy of full-length SEGS-1, 89 bp of upstream sequence, and 543 bp of downstream sequence were amplified, cloned, and sequenced from healthy TME3 and TME14 plants. **(A)** The sequences from 4 independent plants for each genotype were analyzed. The SEG-1 region was compared to the cloned SEGS-1 sequence (Ndunguru et al. 2016), and the flanking regions were compared to the *M. esculenta* v8.1 reference genome (Phytozyme 13). The SNPs were classified as present in all plants (yellow), present in at least one plant from both genotypes (white), present in at least one TME3 plant (green), and present in at least one TME14 plant. **(B)** The number of SNPs in each class and their distributions in the region are shown. SEGS-1 was divided into 4 segments (see Supplementary Figure 1), with only G and F displaying activity (see Figure 6). **(C)** Diagram SEGS-1 junction sequences. The junction is marked by the staggered line. Flanking nucleotides are in black. SEGS-1 nucleotides at the 3’ junction are in blue and SEGS-1 nucleotides at the 5’ junction are in red. Potential base-pairing at the junction is shown. No SNPs were detected in the junction sequences.

**Figure 5.**
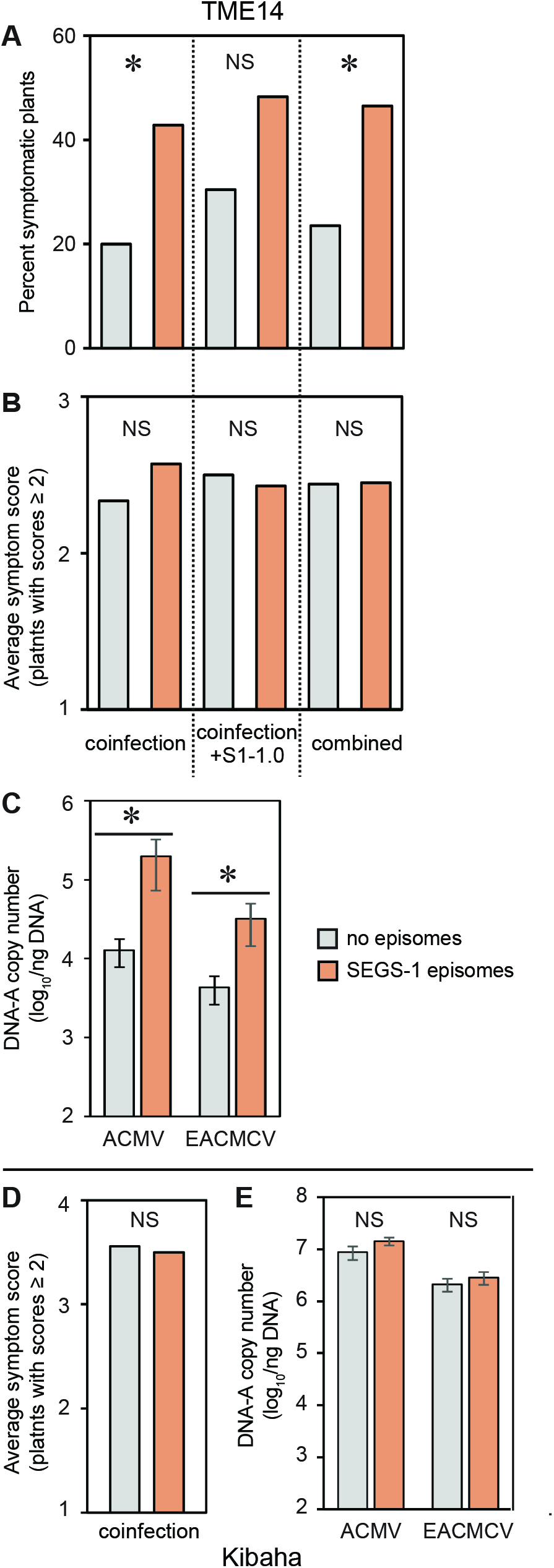
Genomic SEGS-1 episomes increases CMD symptoms and viral copy number in a CMD2 resistant cultivar but in not a tolerant cultivar. **(A)** Incidence of plants (%) with symptom scores ≥2. **(B** and **D)** Average symptom scores of plants with symptom scores ≥2. **(C** and **E)** Average copy number of viral DNA-A per ng total DNA on a log_10_ scale. The bars represent 2 standard errors. Results for TME14 (CMD2 resistant) are shown panels **(A)**, **(B)** and **(C)**. Results for Kibaha (tolerant) are shown in panels **(D)** and **(E)**. The incidence of infection for Kibaha was 100% in all treatments (not shown). Asterisks indicate significant differences (p-value<0.05) between coinfection and coinfection+S1.0 or ± SEG-1 episome treatments within a cultivar. A Mann Whitney test was used in A, B, and D. A two-tailed Students’ T test was used in C and E.

**Figure 6.**
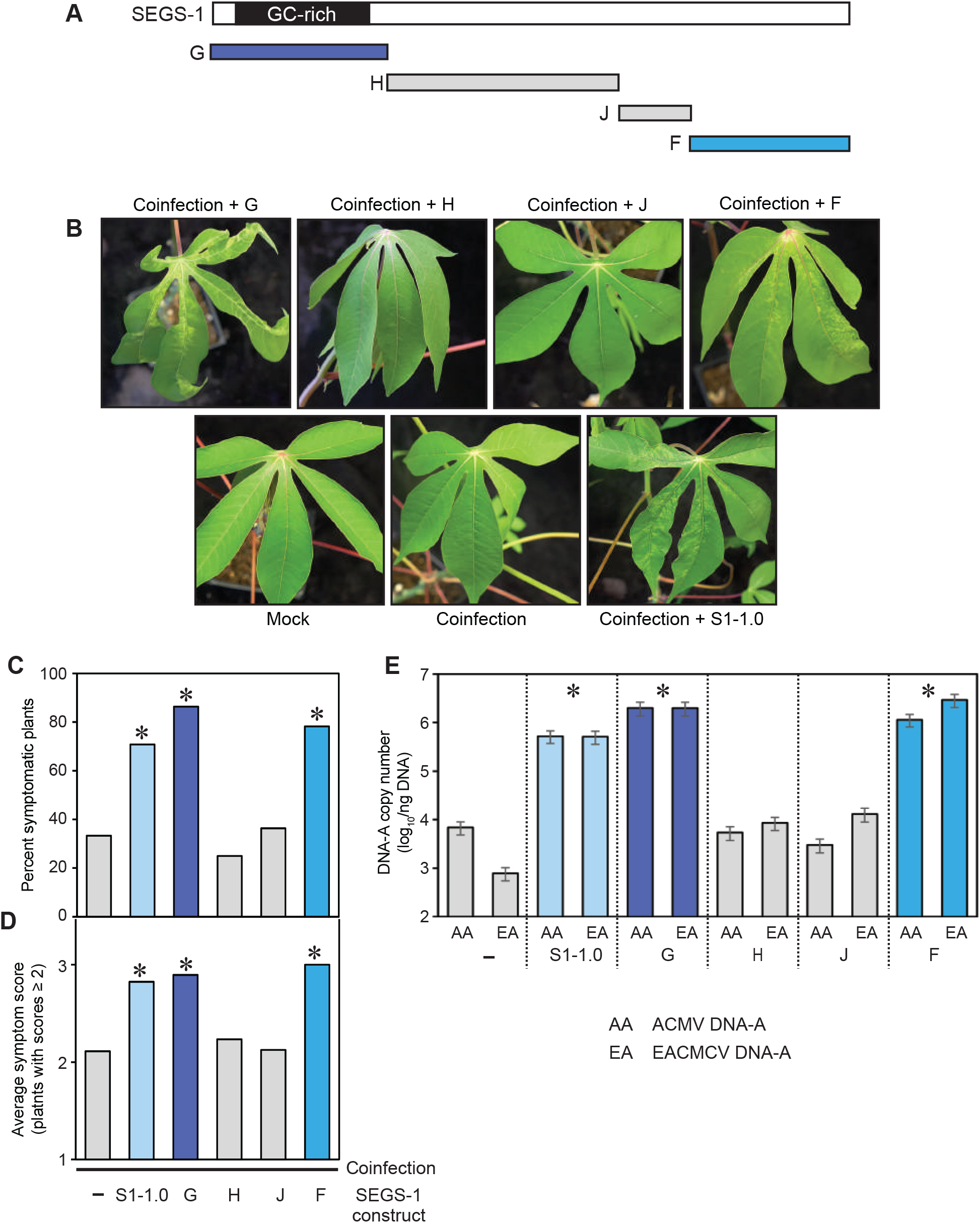
The regions of SEGS-1 that enhance CMB infection in a CMD2 cultivar. **(A)** Exogenous SEGS-1 was cloned into four segments G, H, J and F (see Supplementary Figure 1). **(B)** Images of leaves from TME3 plants co-inoculated with ACMV+EACMCV alone (coinfection) or in combination with exogenous SEGS-1 DNA segments (coinfection + G, H, J or F) under controlled conditions at 28 dpi. The coinfection + S1-1.0 treatment was a positive control **(C)** Incidence of plants (%) with symptom scores ≥2. **(D)** Average symptom severity of plants with symptom scores ≥2. **(E)** Average copy number of viral DNA-A per ng total DNA on a log_10_ scale. The bars represent 2 standard errors. Asterisks indicate significant differences (p-value<0.05) between coinfection and coinfection+SEGS-1 segments treatments. A Mann Whitney test was used in **(C)** and **(D)**, and a two-tailed Students’ T test was used **(E)**.

The presence of SEGS-1 episomes was assessed in total DNA samples from cassava plants. The DNA (100 ng) was amplified by rolling circle amplification (RCA) at 40°C for 2 h using the EquiPhi29 DNA polymerase (ThermoFisher Scientific, USA), as described previously (Rajabu et al., 2023). The RCA product was diluted 10-fold with DNase-free water and 1 μL was used as template in a 50-μL PCR reaction containing the primer pair, S1-4F and S1-2R (Supplementary Table 1), using previously established conditions (Ndunguru et al., 2016; Rajabu et al., 2023).

### Sequencing CMD2 mutations and SEGS-1 genomic copies

The published CMD2 mutations in *MePOLD1* (Manes. 12G077400) in the TME3, TME14, and Kibaha plants (Lim et al., 2022) used in our studies were confirmed by Sanger sequencing. Total DNA was extracted from healthy, young leaves corresponding to the three cassava varieties using the DNeasy Plant Mini Kit (Qiagen). The *M. esculanta* reference genome v8.1 on Phytozyme 13 (https://phytozome-next.jgi.doe.gov) was used to design primer pairs that amplified across the CMD2 SNPs associated with the G680V and A684G amino acid changes (G680VF and GV680R) and with the V528 L replacement (CMD2snEF and CMD2snER) (Supplementary Table 2). The primers were used for end-point PCR to amplify the target regions from plant genomic DNA in a 50-µL reaction containing 100 ng of total DNA, 1.25 U of Hotstart Taq Polymerase (New England Biolabs), 10 µM of each primer, and 1× PCR buffer. PCR conditions were initial denaturation at 95°C for 5 min, 30 cycles at 95°C for 30 s, annealing at 55°C for 30 s, extension at 72°C for 30 s, and final extension at 72°C for 7 min. The PCR products was resolved in 1% agarose gel and the target band was gel purified using a QIAquick Gel Extraction kit (Qiagen, Hilden, Germany). The PCR products were subsequently subjected to direct Sanger sequencing using G680V-F to for the adjacent G680V – A684G SNPs and V528L Seq2 for the V528L SNP. Decoding of the target mutations was done by superimposing the reference sequence with the sequencing chromatograms, followed by manual examination for a heterozygous (double) peaks representing the SNP-specific bases. Validation of the Sanger sequencing results was done by examining mismatches with the reference sequence and the extent of baseline noise in the sequencing chromatogram.

The full-length copies of SEGS-1 and flanking genomic sequences were amplified, cloned, and sequenced from healthy TME3 and TME14 plants. The *M. esculenta* v8.1 reference genome (https://phytozome-next.jgi.doe.gov/) was used to design a primer pair (S1_PyR3 and S1_PyF9; Supplementary Table 2) that amplified a 1639-bp product containing 89 bp of upstream sequence and 543 bp of downstream sequence flanking the 1007-bp SEGS-1 sequence. The presence of 1.5 kb AT-rich region precluded the design of a more distal upstream PCR primer. LongAmp® Hot Start Taq DNA Polymerase kit (New England Biolabs) was used according to the manufacturer’s protocol to amplify the SEGS-1 region from DNA samples collected from 4 plants of each genotype. The PCR products were gel purified and cloned into the pMiniT 2.0 vector using the New England Biolabs PCR Cloning Kit according to the manufacturer’s instructions. Colony PCR using the LongAmp® Hot Start Taq DNA Polymerase and the forward and reverse primers provided in the PCR cloning kit were used to identify positive colonies. Plasmids from positive colonies were purified using QIAprep Spin Miniprep Kit (Qiagen) and subjected to Sanger sequencing using overlapping primers designed based on the *M. esculenta* v8.1 reference genome to obtain coverage of the entire cloned SEGS-1 region. A list of the sequencing primers is found in Supplementary Table 2. Sequences from each plant were aligned to the reference sequence using the SnapGene desktop software (Dotmatics) to assemble a consensus sequence for each full length insert. The resulting consensus sequences were compared to the cloned SEGS-1sequence reported by Ndunguru et al. 2016 and the *M. esculenta* v8.1 reference genome.

## Results

### SEGS-1 does not increase disease severity in susceptible cassava varieties

An earlier study showed that exogenously applied SEGS-1 DNA increases the symptoms caused by ACMV in the hypersusceptible cassava cv. 60444 (Ndunguru et al., 2016). We asked if exogenous SEGS-1 also impacts CMD symptoms and viral DNA copy number in two African cultivars – Namikonga and Kibaha (Amuge et al., 2017; Kulembeka et al., 2012). Both cultivars are highly susceptible coinfection by ACMV and EACMCV, which have been to act synergistically during CMB infection (Fondong et al., 2000). Unlike the earlier studies, we co-inoculated plants with ACMV + EACMCV because there was less plant-to-plant variability in coinfection experiments than in single infection experiments.

Namikonga and Kibaha plants were co-infected with ACMV + EACMCV in the presence and absence of exogenous SEGS-1 DNA under controlled conditions. These experiments used a partial tandem copy of SEGS-1 with duplicated CG-rich regions (S1-1.5a) (Rajabu et al., 2023). Our bombardment inoculation protocol resulted in 100% infection rates in both cultivars, with all inoculated plants showing leaf yellowing and deformation at 28 dpi (Figure 1A). Namikonga and Kibaha had symptom scores ranging from 3 to 4 (on a scale of 1-5) (Supplementary Figure 2). There were no differences in the average symptom scores for either variety whether the plants were co-inoculated with ACMV + EACMCV in the absence or in the presence of exogenous SEGS-1 (Figure 1B). There were also no differences in the DNA copy numbers of ACMV-A and EACMCV-A between the inoculation treatments for both cultivars. These conclusions were supported by within-variety statistical comparisons of the two inoculation treatments, all of which gave p-values > 0.05 in Wilcoxon ranked sum tests for symptom scores and Student’s t tests for viral DNA copy number. Thus, exogenous SEGS-1 did not enhance CMD in Namikonga or Kibaha co-infected with ACMV + EACMCV.

### SEGS-1 alters the CMD response of a CMD2 resistant cultivar

An earlier study (Ndunguru et al., 2016) showed that exogenously applied SEGS-1 DNA can overcome host resistance in the cassava cultivar TME3. We compared the impact of exogenous SEGS-1 on CMD resistance using three resistant varieties, TMS30572, TME14 and TME3 (Table 2). TMS30572 carries the polyploid CMD1 resistance, while TME14 and TME3 are closely related varieties with the CMD2 locus. CMD2 resistance has been correlated with a nonsynonymous, single nucleotide polymorphism in a DNA *polymerase subunit* (*Me*POLD1) (Lim et al., 2022). TME14 and TME3 have the same chimeric mutation that results in a G680V amino acid substitution in POLD1. We confirmed that the TME14 and TME3 plants used in our studies carry the mutation using PCR to amplify the exon 18 of the *MePOLD1* gene from genomic DNA followed by Sanger sequencing (Supplementary Figure 3).

**TABLE 1.** LS means results of viral copy number as a response to SEGS1.

| Source | N | LS Means | Standard Error | P-Value |
| --- | --- | --- | --- | --- |
| Namikonga |  |  |  |  |
| Coinfected | 54 | 1.58 | 0.03 | 0.4485 |
| Coinfected+SEGS1 | 50 | 1.61 | 0.03 |  |
| Kibaha |  |  |  |  |
| Coinfected | 78 | 1.58 | 0.03 | 0.122 |
| Coinfected+SEGS1 | 90 | 1.52 | 0.03 |  |
| TME14 |  |  |  |  |
| Coinfected | 77 | 1.28 | 0.025 | 0.0359 |
| Coinfected+SEGS1 | 62 | 1.35 | 0.025 |  |
| TME3 |  |  |  |  |
| Coinfected | 54 | 1.74 | 0.27 | 0.0002 |
| Coinfected+SEGS1 | 54 | 2.87 | 0.3 |  |
| TMS300572 |  |  |  |  |
| Coinfected | 22 | 1.40 | 0.07 | 0.1041 |
| Coinfected+SEGS1 | 22 | 1.3 | 0.07 |  |
| Mkombozi |  |  |  |  |
| Coinfected | 66 | 1.24 | 0.05 | 0.3151 |
| Coinfected+SEGS1 | 15 | 1.35 | 0.09 |  |

**TABLE 2.**
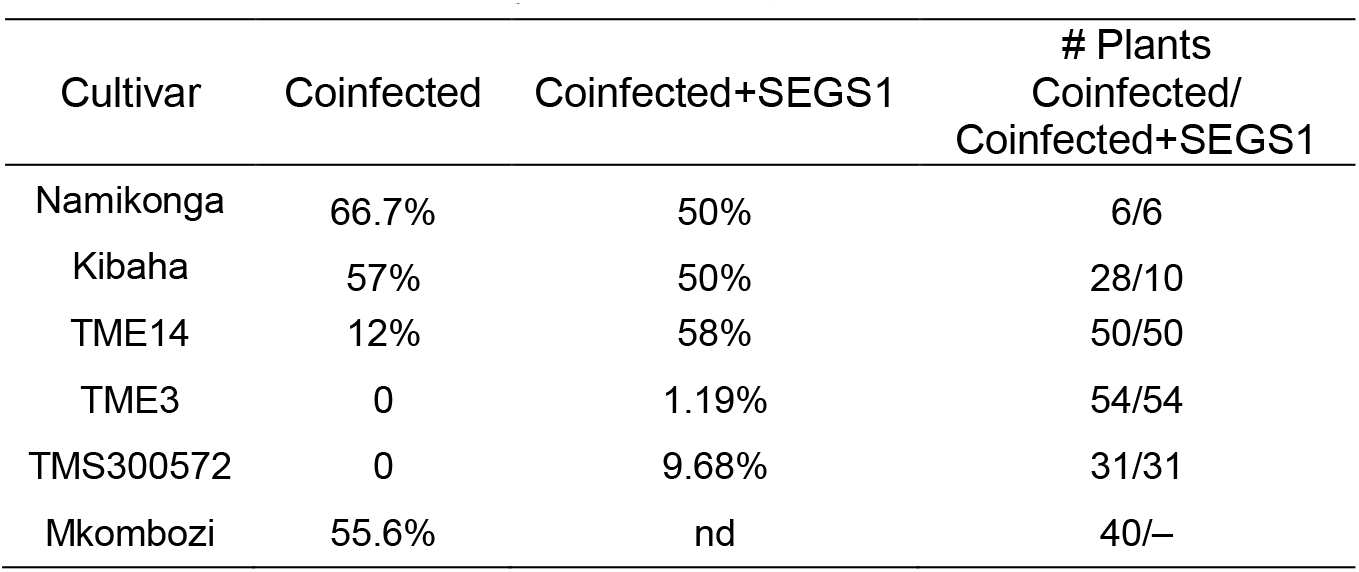
Percent of infected plants with episomes by CMD resistant cultivars.

TME14, TME3, and TMS30572 plants were co-inoculated with ACMV + EACMCV alone or in combination with SEGS-1 DNA (S1-1.5a). Mild symptoms were observed on TME14 and TME3 plants (Figure 2A), with the percent of symptomatic plants approximately 2-fold higher in the presence of SEGS-1 DNA (Figure 2B). Interestingly, the average symptom scores only showed a significant increase for TME3 and not for TME14 (Figure 2C). Both CMD2 cultivars showed significant increases in the DNA copy numbers for ACMV-A and EACMCV-A in the presence of SEGS-1, with TME3 showing a larger increase than TME14 (Figure 2D). In contrast, the presence of SEGS-1 had no detectable effects on the percent of symptomatic plants, average symptom scores, or viral DNA copy numbers in TMS30572 at 46 dpi. (No viral DNA was detected by qPCR in TMS300572 at 28 dpi.)

### Episomal copies of SEGS-1 in cassava varieties

SEGS-1 episomes have been detected in leaves collected from infected plants in Tanzania and Cameroon (Ndunguru et al., 2016). However, it was not possible to assess whether the SEGS-1 episomes in the field-grown plants were from endogenous or exogenous sources. Hence, we asked if SEGS-1 episomes could be detected in infected cassava plants grown under controlled laboratory conditions that precluded potential exogenous sources. For this experiment, cassava plants were co-inoculated with ACMV + EACMCV only. None of the plants were treated with exogenous SEGS-1 plasmid DNA. Total DNA was isolated at 28 dpi and used as template for RCA, which efficiently amplifies small, circular DNA molecules like SEGS-1 episomes. The RCA products were subjected to PCR using a divergent primer pair, (S1-4F/S1-2R) that amplifies an 552-bp product across the SEGS-1 episomal junction but does not amplify the genomic SEGS-1 sequence (Figure 3A) (Ndunguru et al., 2016). Figure 3B shows representative examples of SEGS-1 episomes that were detected in infected Namikonga (lanes 2 and 3), Kibaha (lanes 4 and 5), and TME14 (lanes 6 and 7) plants. No episomes were observed in healthy plants (an example is shown in Figure 3B, lane 1). These results indicated that SEGS-1 episomes are generated during viral infection by a process that most likely involves the full copy of the SEGS-1 sequence in the cassava genome. These results unequivocally established that SEGS-1 episomes are not from exogenous sources like satellite molecules.

We then surveyed the Namikonga, Kibaha, TME14, TME3, and TMS300572 cultivars to gain insight into how often SEGS-1 episomes form when plants are inoculated with ACMV + EACMCV alone or in the presence of exogenous SEGS-1 DNA (S1-1.5a). SEGS-1 episomes were detected in ≥ 50% of Namikonga and Kibaha plants in both inoculation treatments (Table 1). SEGS-1 episomes were also detected in 12% and 58% TME14 plants not treated or treated, respectively, with exogenous SEGS-1 DNA at the time of viral inoculation. The higher frequency of SEGS-1 episomes in TME14 plants in the virus plus exogenous SEGS-1 treatment may be due to residual SEGS-1 plasmid DNA, but we think this is unlikely because systemically infected leaves that emerged after inoculation were sampled at 28 dpi. No SEGS-1 episomes were detected in TME3 or TMS300572 plants co-inoculated with ACMV + EACMCV alone, and only a few plants had episomes in the virus plus exogenous SEGS-1 treatment. These results indicated that the endogenous release of SEGS-1 episomes from the cassava genome is cultivar dependent.

The difference in the capacities of TME3 and TME14 to form SEGS-1 episomes could be due to sequence differences between the full-length SEGS-1 copies in their genomes. To address this possibility, we amplified and sequenced the full-length SEGS-1 copy and 89 bp of upstream sequence and 543 bp of downstream sequence from 4 independent plants for each genotype. Comparison of the 8 sequences uncovered heterogeneity between the sequences from individual plants within a genotype as well as differences between the genotypes (Figure 4A and 4B). Eight SNPs occurred in a subset of TME3 and TME14 sequences (Figure 4A, white lines). The SNPs that were observed in one genotype only occurred in one of the four sequences from that genotype (Figure 5A, green lines – TME3; pink lines – TME14). The 4 SNPs (2 in SEGS-1 and 2 in the distal downstream region) that were conserved across all TME3 sequences were also conserved in all TME14 sequences (Figure 4A, yellow lines). No SNPs occurred around the junction sequences of either genotype (Figure 4C). Thus, it is unlikely that sequence differences in and around the SEGS-1 locus account for the different capacities of TME3 and TME14 to form SEGS-1 episomes.

### Correlation of SEGS-1 episomes with CMD severity

We asked if the presence of SEGS-1 episomes in TME14 correlates with increased frequency of infection, symptom severity and/or viral copy number. When TME14 plants were co-inoculated with ACMV + EACMCV alone or in combination with SEGS-1 DNA (S1-1.5a), the frequency of symptomatic plants was ca. 2-fold higher when SEGS-1 episomes were present (Figure 5A). The copy numbers of ACMV-A and EACMCV-A were also significantly higher (ca. 10-fold) at 28 dpi in plants with SEGS-1 episomes versus plants without episomes (Figure 5C). In contrast, there was no difference in the average symptom scores of plants with or without SEGS-1 episomes (Figure 5B). However, it is important to note that the average is determined using only symptomatic plants (score ≥ 2), and many more plants were symptomatic when SEGS-1 episomes were present (Figure 5A). A similar analysis of Kibaha did not detect any differences in symptom severity (Figure 1B) or viral DNA copy numbers (Figure 1C) between plants with or without SEGS-1 episomes.

### Characterization of sequences required for SEGS-1 activity

TME3 plants do not readily form SEGS-1 episomes (Table 1) but show increased the percent of symptomatic plants, symptom scores, and DNA copy numbers of ACMV-A and EACMCV-A when treated with exogenous SEGS-1 DNA (Figure 2). These properties allowed us to characterize the sequence requirements for SEGS-1 activity in the absence of endogenous background activity that could mask the effects of exogenously applied SEGS-1 sequences. SEGS- 1 was subcloned into 4 fragments, designated as SEGS-1 regions G (positions 1–277), I (positions 275–646), J (positions 644–756), and F (positions 757–1007) (Figure 6A and Supplementary Figure 1). Each cloned fragment was co-inoculated with ACMV + EACMCV onto TME3 plants. The positive control was co-inoculation of both viruses with the full-length SEGS-1 sequence, while the negative control was plants only inoculated with both viruses. The percent symptomatic plants, average symptom scores, and viral DNA-A copy numbers were assessed at 28 dpi for all inoculation treatments. Virus-inoculated plants treated with fragments G or F developed clear symptoms that were similar to the full-length SEGS-1 positive control (S1-1.0), while plants treated with fragments J and H showed few symptoms like the virus only negative control (Figure 6B). The percent of infected plants were similar for plants treated with S1-1.0, G or F and ca. 2- fold higher than the virus alone control, or plants treated with J or H (Figure 5C). The S1-1.0, G, and F treatments also resulted in more severe symptoms (Figure 6D) and >10-fold increases in viral DNA-A copy numbers (Figure 6E) than the virus alone, H and J treatments. These differences were statistically significant (p-values < 0.05). These results established that the G and F regions are separately sufficient for SEGS-1 activity, and that the intervening J and H regions are not required for activity.

Fragments G and F are at the opposite ends of the full-length copy of SEGS-1 in the cassava genome but are adjacent in the episome and flank the junction sequence. We fused G and F to reconstitute the junction region in fragment N (Supplementary Figure 1). Co-inoculation of the cloned fragment N with ACMV + EACMCV also resulted in statistically significant increases in the percent of symptomatic plants, symptom scores, and viral DNA-A copy numbers when compared to virus alone plants (Figure 7B, 7C, and 7D). However, no differences were detected when the frequency of infected plants, symptom severity, and viral DNA copy numbers were compared between the F, G and N treatments, indicating that joining the F and G regions and reconstituting the junction sequence does not result in increased SEGS-1 activity.

**Figure 7.**
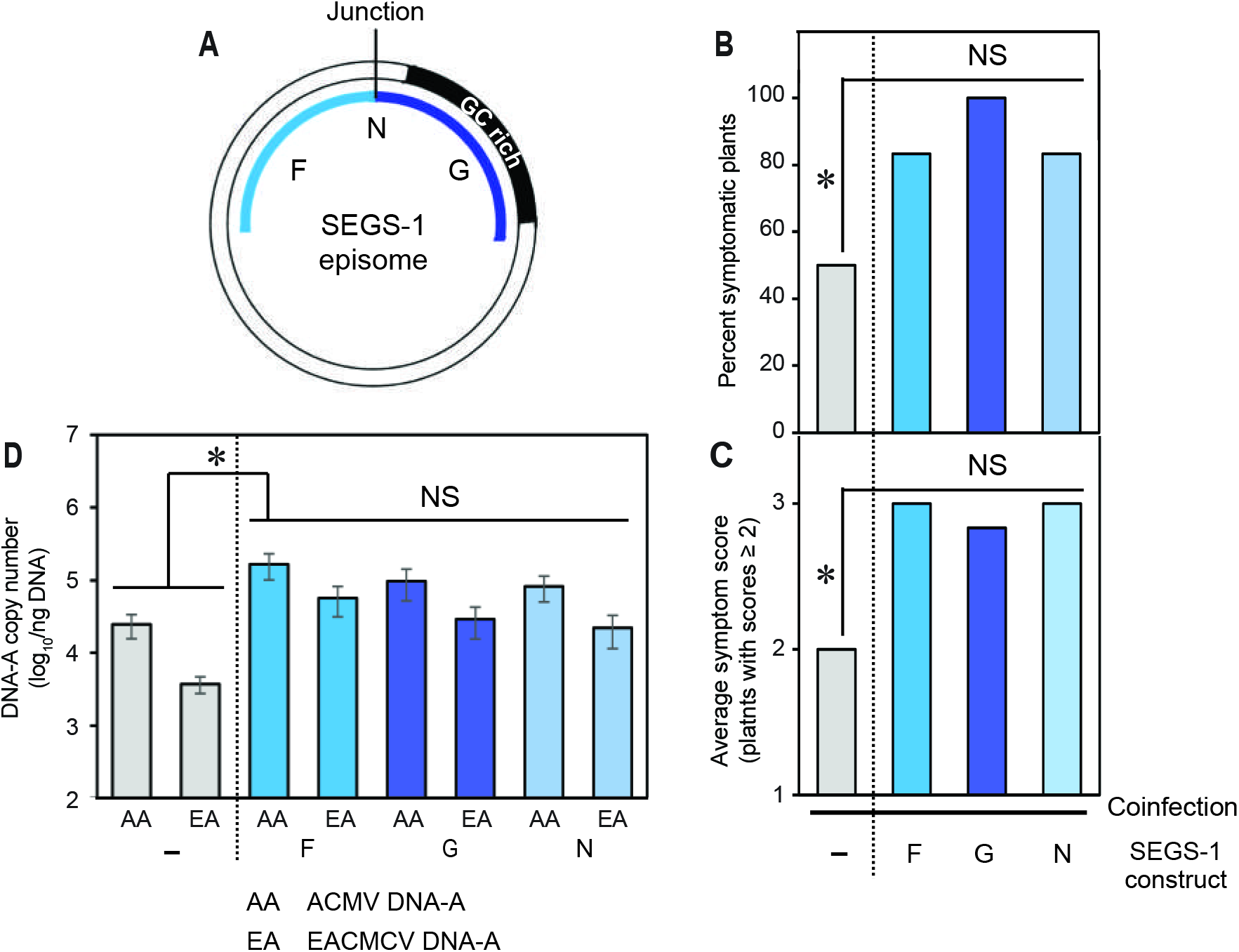
The SEGS-1 episome junction does not increase activity. **(A)** Diagram of the SEGS-1 episome showing the N region that combines the F and G regions to reconstitute episome junction sequences. **(B)** Incidence of plants (%) with symptom scores ≥2. **(C)** Average symptom severity of plants with symptom scores ≥2. **(D)** Average copy number of viral DNA-A per ng total DNA on a log_10_ scale. The bars represent 2 standard errors. Asterisks indicate significant differences (p-value<0.05) between coinfection and coinfection+SEGS-1 segments treatments. No differences were detected between the SEGS-1 regions. A Mann Whitney test was used in **(B)** and **(C)**, and a two-tailed Students’ T test was used **(D)**.

## Discussion

SEGS-1 was first identified as a cassava genomic sequence that also occurs as an episomal DNA in cassava plants showing severe CMD symptoms in the field (Ndunguru et al., 2016). A previous study showed that SEGS-1 can increase disease severity in cassava plants co-inoculated with a CMB and exogenous SEGS-1 DNA (Ndunguru et al., 2016). We report here that the effects of exogenous SEGS-1 on CMD varies between cassava cultivars, with CMD2 cultivars most affected. We also show that cultivars differ in their capacities to produce endogenous SEGS-1 episomes from the full-length genomic copy of SEGS-1, and that there is a correlation between the generation of endogenous SEGS-1 episomes and disease severity in a CMD2 cultivar. Taken together, these results established that the host genetic background is an important determinant of SEGS-1 activity during CMB infection in cassava.

In our studies, CMD2 resistant cultivars but not the susceptible cultivars developed more severe disease when ACMV and EACMCV were co-inoculated with exogenous SEGS-1 DNA. Disease severity was assessed at 28 dpi using three parameters – the percent of symptomatic plants, the average symptom score, and viral DNA copy number. The parameter values were high in susceptible plants (Namikonga and Kibaha) in the absence of exogenous SEGS-1 DNA and may have masked any effects of SEG-1. For example, 100% of the plants were symptomatic in the absence and the presence of exogenous SEGS-1 DNA, precluding the detection of a change in the parameter. We reported previously that exogenous SEGS-1 DNA increases symptom severity in the hypersusceptible cassava cv. 60444 during single infections with ACMV, EACMCV or East African cassava mosaic virus-Uganda (EACMV-Ug) (Ndunguru et al., 2016). A single virus infection with a CMB is often milder than a coinfection with two CMBs (Fondong et al., 2000), and the effect of SEGS-1 may be more readily detected in a single infection. It is unlikely that the synergistic effects of an ACMV+EACMCV coinfection suppress SEGS-1 activity because exogenous SEGS-1 can increase disease severity during coinfection of CMD2 cultivars (Vanitharani et al., 2004). The CMD2 cultivars have lower disease parameter values in the absence of SEGS-1 that are comparable to single-virus infections. Thus, we think it is likely the high disease parameter values of susceptible cultivar coinfections preclude detection of SEGS-1 activity.

Co-inoculation of exogenous SEGS-1 plasmid DNA with EACMV-Ug resulted in severe symptoms in TME3, a CMD2 resistant cassava cultivar (Ndunguru et al., 2016). In this study, we found that exogenous SEGS-1 increased the percentage of infected plants and viral DNA copy number significantly in two CMD2 resistant cultivars (TME14 and TME3) but not in a CMD1 resistant cultivar (TMS300572). The resistance phenotypes of CMD1 and CMD2 plants have different genetic origins that may determine how SEGS-1 effects CMD. CMD1 resistance is polygenic and recessive (Okogbenin et al., 2013), and the genes involved in the resistance are not known. CMD2 resistance is mediated by a single, dominant gene encoding DNA polymerase subunit 1 (*MePOLD1*) (Akano et al., 2002; Lim et al., 2022). TME3 and TME14 carry the same *MePOLD1* amino acid mutation (G680V in exon 18), and they have similar responses to exogenous SEGS-1. Interestingly, Kibaha is also a *MePOLD1* variant but has a different amino acid polymorphism (A684G in exon 13) that does not confer resistance in our CMB coinfection studies.

SEGS-1 episomes were first detected in field grown plants with CMD (Ndunguru et al., 2016). We report here the presence of SEGS-1 episomes in cassava plants coinfected with ACMV and EACMCV under laboratory conditions. Importantly, the detection of SEGS-1 episomes did not depend on treatment of the infected plants with exogenous SEGS-1 DNA. The only source of the episomes in untreated plants is the full-length copy of SEGS-1 that occurs in all characterized cassava genomes. Although the genomic sequence of SEGS-1 is highly conserved, cultivars differ in their capacities to form SEGS-1 episomes during CMD. More than 50% of infected Namikonga and Kibaha plants formed episomes. Kibaha plants with endogenous SEGS-1 episomes did not have higher symptom scores or viral titers than infected plants lacking episomes, supporting the idea that SEGS-1 effects are not readily detected in highly susceptible cultivars. Endogenous SEGS-1 episomes were also detected the CMD2 resistant cultivar TME14 during coinfection, with 14% of the plants containing episomes. Unlike the susceptible cultivars, the presence of SEGS-1 episomes in TME14 correlated with higher incidence of infection and viral titers, suggesting that that the formation of endogenous SEGS-1 episomes causes more severe disease in a CMD resistant cultivar. This effect is cultivar dependent because no endogenous SEGS-1 episomes were found in TME3 or TMS300572.

The failure of TME3 to generate endogenous episomes was unexpected because of its close relationship to TME14 (Bredeson et al., 2016). However, hierarchical clustering of residual genetic distances has indicated that TME3 and TME14 genomes are closely related but not identical (Rabbi et al., 2014), supporting the idea that there is something different between their genomes that impacts the generation of SEGS-1 episomes. Sequence comparison of the genomic copies of SEGS-1 and flanking regions did not uncover any differences in TME3 and TME14 that are likely to reflect this genetic difference. Thus, we hypothesize that the different capacities of TME3 and TME14 to form endogenous SEGS-1 episomes is due to a SNP elsewhere in the cassava genome. Accordingly, TME14 and other episome-forming cultivars may encode a protein that acts in trans to generate SEGS-1 episomes, and that the TME3 genome contains a mutation that interferes with its production or activity. This idea is consistent with the observation that the longest open reading frame in SEGS-1 only specifies a 48 amino acid peptide (ORF positions 164–18), lending further support to our premise that episome release depends on a protein encoded at another location in the genome. A similar scenario occurs for transposable elements, which are often defective and depend on transposase genes encoded distally in a plant genome for their mobilization (Pulido & Casacuberta, 2023).

A SEGS-1 transgene increases ACMV symptom severity in Arabidopsis, establishing that SEGS-1 by itself is sufficient to enhance begomovirus infection (Rajabu et al., 2023). This observation provides an explanation for why exogenous SEGS-1 increases disease severity in TME3. It also raises the question as to whether SEGS-1 episomes are necessary for disease enhancement in cassava or if the SEGS-1 genomic copy is sufficient. The strongest evidence supporting the involvement of episomes comes from TME14, which showed a positive correlation between episome presence and disease severity. Moreover, the SEGS-1 genomic sequences of TME3 and TME14 are essentially identical and should impact CMD similarly if the genomic copy is the active form. Given that SEGS-1 belongs to a repetitive sequence family, it is likely located in a heterochromatic region of the genome that is functionally inactive. SEGS-1 episomes are only found in CMB-infected plants, consistent with infection promoting episome formation. Begomovirus infection induces host DNA replication (Ascencio-Ibánez et al., 2008; Nagar et al., 2002) and interferes with DNA methylation (Gui et al., 2022). The opening of host chromatin during replication and/or the decrease in DNA methylation could lead to activation of the SEGS- 1 genomic copy or the generation of SEGS-1 episomes.

The strong effect of exogenous SEGS-1 DNA on CMB infection in TME3 argues that its genomic copy does not enhance CMD. Instead, SEGS-1 activity in cassava is likely mediated by extrachromosomal SEGS-1 DNA through the formation of endogenous episomes or the application of exogenous SEGS-1 plasmid DNA. Endogenous SEGS-1 episomes have some features in common with but are distinct from extrachromosomal circular DNAs (eccDNAs) that have been characterized in plants (Peng et al., 2022; Zhuang et al., 2024). Like SEGS-1, the generation of some eccDNAs has been associated with stress and epigenetic changes. Conversely, SEGS-1 episomes are larger than most eccDNAs and correspond to a specific cassava sequence with a conserved junction, while eccDNAs appear to be generated randomly from the plant genome and show little reproducibility or sequence conservation with each other.

We mapped SEGS-1 activity to two regions, F and G that flank the episomal junction. Both regions increased CMD symptoms and viral DNA accumulation when applied separately to TME3. When F and G were combined to restore the episomal junction in N, no additional disease enhancement was observed. The lack of synergy between F and G is consistent with both regions of SEGS-1 targeting the same host defense pathway. Moreover, the episomal junction sequences do not contribute directly to SEGS-1 disease enhancement. The SEGS-1 episomal junction is imbedded in a 10-bp repeat motif with a single mismatch at the junction site that may be involved in SEGS-1 episome formation. Thus, we hypothesize that SEGS-1 disease enhancement involves two separable steps – [1] episomal formation involving the junction sequences and a trans acting protein encoded elsewhere in the genome and [2] disease enhancement mediated the F and/or G regions. The SEGS-1 products and how they function is not known. Unpublished results in Arabidopsis showed that the G region, which includes the longest open reading frame, is transcribed from a SEGS-1 transgene (Ruhel et al., in preparation). The G region overlaps 15 SEGS-1 related sequences that are distributed across 10 chromosomes of the cassava genome, while the F region overlaps 2 SEGS-1 related sequences that occur on separate chromosomes (Ndunguru et al., 2016). It is not known if any of the SEGS-1 related sequences occur in episomes or can enhance CMD severity, but several have low E values and high identities in BLAST comparisons to the full-length genomic copy of SEGS-1, raising the possibility that they are also active during CMD.

The studies reported here describe a new type of plant virus/host interaction in which viral infection leads to the generation of DNA episomes from SEGS-1 genomic DNA and an increase in CMD severity. The SEG-1 genomic sequence is ubiquitous across cassava cultivars, and 3 of the 5 cultivars in this report generate SEGS-1 episomes during CMB infection. Thus, the endogenous SEGS-1/virus interaction is likely to be a common feature of the CMB infection process that may be reflected by the severe CMD observed in some African fields planted with CMD-resistant cassava. This is most concerning for CMD2-resistant plants, which typically become infected and then undergo recovery. CMD2 cultivars that produce SEGS-1 episomes during the infection phase are likely to be less able undergo recovery and, instead, develop severe CMD. Hence, it is important to identify cultivars like TME14 that produce SEGS-1 episomes and remove them from cassava breeding programs, if alternative cultivars like TME3 that do not generate episomes are available. As information becomes available about cultivar-specific generation of SEGS-1 episomes, its inclusion into cassava genetic databases could be used to identify molecular markers associated with episome formation. Such markers could be used to identify new cultivars with reduced risk of episome formation and potential gene editing targets to prevent episome formation, as well as provide insight into the mechanisms underlying SEGS-1 episomes formation and mode(s) of action. An integrated approach involving virology, genetics, and breeding to minimize the effects of SEGS-1 episomes will be essential to develop effective and sustainable CMD resistant cassava.

## Acknowledgments

This study was funded by a grant # OPP1149990 to L.H.-B., J.N. and J.A.I. from the Bill & Melinda Gates Foundation.

---

**Supplementary Figure 1.** SEGS-1 fragments. **(Top)** Properties of SEGS-1 fragments. In the primer sequences, black marks SEGS-1 sequences, red marks the *Not*I linker, and blue marks added residues not in the linker or SEGS-1. **(Bottom)** The sequences of the SEGS-1 fragments. The green shading indicates the GC-rich region. The blue line marks the junction position. The underlined nucleotides overlap the adjacent fragment.

**Supplementary Figure 2.** CMD symptom scoring. Images of leaves illustrating the symptom scores. Description of the extent of symptoms is below each image.

**Supplementary Figure 3.** Confirmation of CMD2 mutations. Sanger sequencing profiles for the TME3, TME14, Kibaha, and TMS30752 plants used in our studies. The codons in exon 18 of the *MePOLD1* gene (Manes. 12G077400) containing the nonsynonymous SNP haplotypes, G680V and A684G, is shown at the top. The G680V and A684G SNPs are highlighted in red in each profile. Except for TMS30752 used as control negative, The TME3 and TM14 chromatograms show double peaks in codon 680 indicating that they are chimeric for wild-type GGT (glycine) and mutant GTT (valine) at this position. The Kibaha chromatogram shows a double peak in codon 684 indicating that it is chimeric for wild-type GCC (alanine) and mutant GGC (glycine) at this position. TMS30752, which is not a CMD2 cultivar, does not have a mutation at either position. Codons 680 and 684 are marked and the SNPs are indicated by an asterisk (*).

## References

1. Aimone, C., De León, L., Dallas, M., Ndunguru, J., Ascencio-Ibáñez, J., & Hanley-Bowdoin, L. (2021a). A new type of satellite associated with cassava mosaic begomoviruses. Journal of Virology, 95(21), 10.1128/jvi.00432-00421.

2. Aimone, C., Hoyer, J. S., Dye, A. E., Deppong, D. O., Duffy, S., Carbone, I., & Hanley-Bowdoin, L. (2022). An experimental strategy for preparing circular ssDNA virus genomes for next-generation sequencing. Journal of Virological Methods, 300, 114405.

3. Aimone, C., Lavington, E., Hoyer, J. S., Deppong, D. O., Mickelson-Young, L., Jacobson, A., … Duffy, S. (2021b). Population diversity of cassava mosaic begomoviruses increases over the course of serial vegetative propagation. Journal of General Virology, 102(7), 001622.

4. Akano, A., Dixon, A., Mba, C., Barrera, E., & Fregene, M. (2002). Genetic mapping of a dominant gene conferring resistance to cassava mosaic disease. Theoretical and Applied Genetics, 105, 521–525.

5. Amuge, T., Berger, D. K., Katari, M., Myburg, A. A., Goldman, S., & Ferguson, M. E. (2017). A time series transcriptome analysis of cassava (Manihot esculenta Crantz) varieties challenged with Ugandan cassava brown streak virus. Scientific reports, 7(1), 9747.

6. Ascencio-Ibáñez, J., Dallas, M., & Hanley-Bowdoin, L. (2023). Begomovirus Inoculation in Arabidopsis and Cassava. In Plant-Virus Interactions (pp. 71–79): Springer.

7. Ascencio-Ibánez, J. T., Sozzani, R., Lee, T.-J., Chu, T.-M., Wolfinger, R. D., Cella, R., & Hanley-Bowdoin, L. (2008). Global analysis of Arabidopsis gene expression uncovers a complex array of changes impacting pathogen response and cell cycle during geminivirus infection. Plant physiology, 148(1), 436–454.

8. Bayata, A. (2019). Review on nutritional value of cassava for use as a staple food. Sci J Anal Chem, 7(4), 83–91.

9. Bredeson, J. V., Lyons, J. B., Prochnik, S. E., Wu, G. A., Ha, C. M., Edsinger-Gonzales, E., … Egesi, C. (2016). Sequencing wild and cultivated cassava and related species reveals extensive interspecific hybridization and genetic diversity. Nature biotechnology, 34(5), 562–570.

10. Crespo-Bellido, A., Hoyer, J., Dubey, D., Jeannot, R., & Duffy, S. (2021). Interspecies recombination has driven the macroevolution of cassava mosaic begomoviruses. Journal of Virology, 95(17), 10.1128/jvi.00541-00521.

11. Deng, D., Otim-Nape, W., Sangare, A., Ogwal, S., Beachy, R., & Fauquet, C. (1997). Presence of a new virus closely related to East African cassava mosaic geminivirus, associated with cassava mosaic outbreak in Uganda.

12. Duffy, S., & Holmes, E. (2009). Validation of high rates of nucleotide substitution in geminiviruses: phylogenetic evidence from East African cassava mosaic viruses. Journal of General Virology, 90(6), 1539–1547.

13. FAOSTAT. (2021). Food and Agriculture Organization of the United Nations statistical database. Available online: fao.org/faostat/en/#data/QCL (accessed March, 2024).

14. Fondong, V., Pita, J., Rey, M., De Kochko, A., Beachy, R., & Fauquet, C. (2000). Evidence of synergism between African cassava mosaic virus and a new double-recombinant geminivirus infecting cassava in Cameroon. Journal of General Virology, 81(1), 287–297.

15. Gong, P., Tan, H., Zhao, S., Li, H., Liu, H., Ma, Y., … Lozano-Durán, R. (2021). Geminiviruses encode additional small proteins with specific subcellular localizations and virulence function. Nature Communications, 12(1), 4278.

16. Gui, X., Liu, C., Qi, Y., & Zhou, X. (2022). Geminiviruses employ host DNA glycosylases to subvert DNA methylation-mediated defense. Nature Communications, 13(1), 575.

17. Hanley-Bowdoin, L., Bejarano, E., Robertson, D., & Mansoor, S. (2013). Geminiviruses: masters at redirecting and reprogramming plant processes. Nature Reviews Microbiology, 11(11), 777–788.

18. Hoyer, J. S., Fondong, V. N., Dallas, M. M., Aimone, C. D., Deppong, D. O., Duffy, S., & Hanley-Bowdoin, L. (2020). Deeply sequenced infectious clones of key cassava begomovirus isolates from Cameroon. Microbiology Resource Announcements, 9(46), 10.1128/mra.00802-00820.

19. Kulembeka, H., Ferguson, M., Herselman, L., Kanju, E., Mkamilo, G., Masumba, E., … Labuschagne, M. T.. (2012). Diallel analysis of field resistance to brown streak disease in cassava (Manihot esculenta Crantz) landraces from Tanzania. Euphytica, 187, 277–288.

20. Legg, J., Jeremiah, S., Obiero, H., Maruthi, M., Ndyetabula, I., Okao-Okuja, G., … Gashaka, G. (2011). Comparing the regional epidemiology of the cassava mosaic and cassava brown streak virus pandemics in Africa. Virus research, 159(2), 161–170.

21. Legg, J., Kumar, P., Makeshkumar, T., Tripathi, L., Ferguson, M., Kanju, E., … Cuellar, W. (2015). Cassava virus diseases: biology, epidemiology, and management. In Advances in virus research (Vol. 91, pp. 85–142): Elsevier.

22. Lim, Y.-W., Mansfeld, B. N., Schläpfer, P., Gilbert, K. B., Narayanan, N. N., Qi, W., … Gehan, J. (2022). Mutations in DNA polymerase δ subunit 1 co-segregate with CMD2-type resistance to Cassava Mosaic Geminiviruses. Nature Communications, 13(1), 3933.

23. Nagar, S., Hanley-Bowdoin, L., & Robertson, D. (2002). Host DNA replication is induced by geminivirus infection of differentiated plant cells. The Plant Cell, 14(12), 2995–3007.

24. Ndunguru, J., De León, L., Doyle, C., Sseruwagi, P., Plata, G., Legg, J., … Ascencio-Ibáñez, J. (2016). Two novel DNAs that enhance symptoms and overcome CMD2 resistance to cassava mosaic disease. Journal of Virology, 90(8), 4160–4173.

25. Okogbenin, E., Setter, T., Ferguson, M., Mutegi, R., Ceballos, H., Olasanmi, B., & Fregene, M. (2013). Phenotypic approaches to drought in cassava. Frontiers in physiology, 4, 93.

26. Peng, H., Mirouze, M., & Bucher, E. (2022). Extrachromosomal circular DNA: a neglected nucleic acid molecule in plants. Current Opinion in Plant Biology, 69, 102263.

27. Pita, J., Fondong, V., Sangare, A., Otim-Nape, G., Ogwal, S., & Fauquet, C. (2001). Recombination, pseudorecombination and synergism of geminiviruses are determinant keys to the epidemic of severe cassava mosaic disease in Uganda. Journal of General Virology, 82(3), 655–665.

28. Pulido, M., & Casacuberta, J. M. (2023). Transposable element evolution in plant genome ecosystems. Current Opinion in Plant Biology, 75, 102418.

29. Rabbi, I., Hamblin, M., Kumar, P., Gedil, M., Ikpan, A., Jannink, J., & Kulakow, P. (2014). High-resolution mapping of resistance to cassava mosaic geminiviruses in cassava using genotyping-by-sequencing and its implications for breeding. Virus research, 186, 87–96.

30. Rajabu, C. A., Dallas, M. M., Chiunga, E., De León, L., Ateka, E. M., Tairo, F., … Hanley-Bowdoin, L. (2023). SEGS-1 a cassava genomic sequence increases the severity of African cassava mosaic virus infection in Arabidopsis thaliana. Frontiers in Plant Science, 14, 1250105.

31. Stanley, J., & Gay, M. R. (1983). Nucleotide sequence of cassava latent virus DNA. Nature, 301(5897), 260–262.

32. Vanitharani, R., Chellappan, P., Pita, J. S., & Fauquet, C. M. (2004). Differential roles of AC2 and AC4 of cassava geminiviruses in mediating synergism and suppression of posttranscriptional gene silencing. Journal of Virology, 78(17), 9487–9498.

33. Yang, X., Wang, B., Li, F., Yang, Q., & Zhou, X. (2016). Research advances in geminiviruses. Current Research Topics in Plant Virology, 251–269.

34. Zhou, X., Liu, Y., Calvert, L., Munoz, C., Otim-Nape, G., Robinson, D., & Harrison, B. (1997). Evidence that DNA-A of a geminivirus associated with severe cassava mosaic disease in Uganda has arisen by interspecific recombination. Journal of General Virology, 78(8), 2101–2111.

35. Zhuang, J., Zhang, Y., Zhou, C., Fan, D., Huang, T., Feng, Q., … Han, B. (2024). Dynamics of extrachromosomal circular DNA in rice. Nature Communications, 15(1), 2413.

