## Supplemental Figures for "SEGS-1 episomes generated during cassava mosaic disease influence disease severity"

Supplementary Figure 1

| Clone name | SEGS-1 fragment | SEGS-1 coordinates | Primer name | Primer sequence | Annealing (°C) |
|---|---|---|---|---|---|
| pNSB2163 | G | 1–277 | SEG1FWDNot1 | GAATGCGGCCGCCACTACGCTACGCAGCAGCC | 63 |
| | | | S1GnnRNot1 | GAATGCGGCCGCAGAGGAGCGAGGTGGGTG | |
| pNSB2166 | H | 275–646 | SATII FwdNot1 | GAATGCGGCCGCGCAGTTCAGCAGTTCAACTG | 48 |
| | | | SATII RNot1 | GAATGCGGCCGCGCTTTTATTAAAAAATTTTGG | |
| pNSB2165 | J | 644–756 | S1InnFwdNot1 | GAATGCGGCCGCTTTACAAAGCTCAGCTTGG | 55 |
| | | | SATII 1RevNot1 | GAATGCGGCCGCTGAAGGATTAGAGGCTACCC | |
| pNSB2164 | F | 757–1007 | S1FnnFwdNot1 | GAATGCGGCCGCACTCTATTTTTCCGTTTGG | 52 |
| | | | SEGS1REVNot1 | GAATGCGGCCGCGTACCACTACGCTACGCAG | |
| pNSB2184 | N | 757–1007 + 1–277 | SEGS1NIFwd | GAATGCGGCCGC ACTCTATTTTTCCGTTTGG | 58 |
| | | | SEGS1NrRev | GAATGCGGCCGCAGAGGAGCGAGGTGGGTG | |

### SEGS-1 Fragments

**G**

ACGCTACGCAG|CAGCCATCATCGACATCGTATTTTAA*CCAGAGGACCCGTCGACCGCCTGAGCAGCAGCACGTC*
*GCACCAGCACCACCGCCGCATCGCGCGCCTGTGAGCCGCCGCACCACTGGATCTCGTGCTCGTGAGCCGCCG*
*CACGCCGCAACTCTTCATCTACCGCTCGTTTACAGCCCACCTCTGTATCACGCGATTGTGAGCCGCCGACTGCCC*
*GCCGCACGCCCGCACCTCTGCATCAACTGCTCGTTTGCCACCCACCTCGCTCCTCT*

**J**

*CTGC*AGTTCAGCAGTTCAACTGTAAGCATTTTTTCGTTAAATCTGAAGAAAATAGTTCTGGATAGAATTTTGATTGGT
AAGCATTATGAATTTATTATGACATTCAAGTTTATAGGCATCATAGTGTTGCTTAGGACATACTTAGCTTGTAGTTCCA
GAAAATAGAGTCATTTCTGGTTTTCTTTTACAATGGAGGTGTTTATTCCATTGTAATTTTGAGCTGAGCTTTGTTAAG
GACCTTTGGAGCTCGAGCTTTGTTTACAAGGCATCTTGATAGAGCTTTTCGAGCTCGAATTAGAATTAGGCTCATG
GTTATACTAAAGGGAGTTTTTCATGAGTTTGAGTGCTTCCAAAATTTTTTAATAAAAGC

**H**

GCTTTACAAAGCTCAGCTTGGATCGATTACACCTCTACTGACCCTACTCAGTTTGGGACTCTGGCTGGGGCCATT
CTCAAAAGCCATTTATCTGGGTAGCCTCTAATCCTTCA

**F**

ACTCTATTTTTCCGTTTGGTTCTGAGAGAGTACTAAAAAGGAAATCCAACCATATATGATCAAATCTAATGATATAGCT
GGTGAGTACTGCAACATAATTGCAATTTATGCAGTTATTTCTCTTGAATTTGGTATCTGCAATTTATGTATAAATCCCT
AGCAGAATATTTTACTGGAGTGGTGAATATGTGTAGGCTTCACTATGGTGGAAATGGAAATTTGTGTGTGATAACTT
CCTAACTGGCTGCT|GC

**N**

ACTCTATTTTTCCGTTTGGTTCTGAGAGAGTACTAAAAAGGAAATCCAACCATATATGATCAAATCTAATGATATAGCT
GGTGAGTACTGCAACATAATTGCAATTTATGCAGTTATTTCTCTTGAATTTGGTATCTGCAATTTATGTATAAATCCCT
AGCAGAATATTTTACTGGAGTGGTGAATATGTGTAGGCTTCACTATGGTGGAAATGGAAATTTGTGTGTGATAACTT
CCTAACTGGCTGCT|CAGCCATCATCGACATCGTATTTTAA*CCAGAGGACCCGTCGACCGCCTGAGCAGCAGCAC*
*GTCGCACCAGCACCACCGCCGCATCGCGCGCCTGTGAGCCGCCGCACCACTGGATCTCGTGCTCGTGAGCCG*
*CCGCACGCCGCAACTCTTCATCTACCGCTCGTTTACAGCCCACCTCTGTATCACGCGATTGTGAGCCGCCGACTG*
*CCCGCCGCACGCCCGCACCTCTGCATCAACTGCTCGTTTGCCACCCACCTCGCTCCTCT*

Supplementary Figure 2

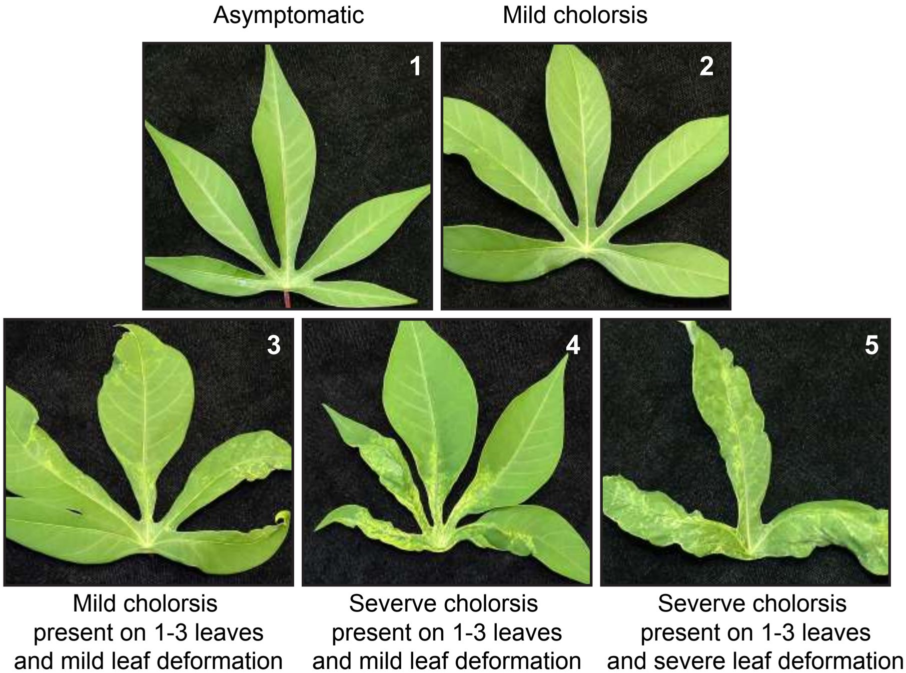

Supplementary Figure 3

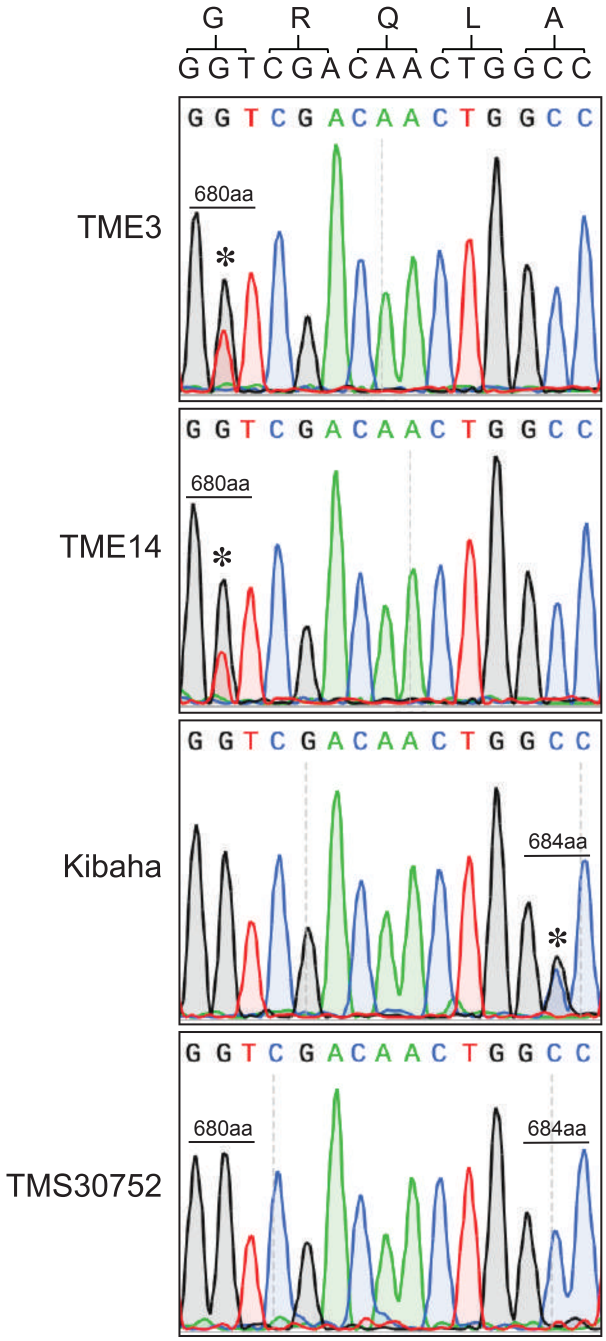
