## Supplemental Table 1 for "SEGS-1 episomes generated during cassava mosaic disease influence disease severity"

| Supplementary Table 1. qPCR and SEGS-1 Episomes Primer Sequences | | | |
| --- | --- | --- | --- |
|  | **Primer** | **Sequence (5'-3')** | **Use** |
| ACMV DNA-A* | P3P-AA2F | TCTGCAATCCAGGACCTACC | qPCR |
|  | P3P-AA2R+4R | GGCTCGCTTCTTGAATTGTC |  |
| EACMCV DNA-A* | EACMVQ1 | GTACCATGCGTCGTTTGAATA | qPCR |
|  | EACMVQ2 | GCAAGTCCCAGAGGAAATAGA |  |
| SEGS-1^Ɨ^ | S1-4F* | GGGTAGCCTCTAATCCTTCA | Episome detection |
|  | S1-2R* | CAGTTGAACTGCTGAACTGC |  |
| *Aimone et al., 2022 | | | |
| ^Ɨ^Ndunguru et al., 2016 | | | |
