## Supplemental Table 2 for "SEGS-1 episomes generated during cassava mosaic disease influence disease severity"

Supplemental Table 2. Primers for cloning and sequencing cassava genomic sequences

| Primer name | Sequence (5’-3’) | Annealing (°C) | Product (bp) |  |
| --- | --- | --- | --- | --- |
| Amplification and cloning of SEGS-1 and flanking genomic regions | | |  |  |
| S1_PyF9 | TGCGTAGTTTGGTGTGTTACT | 55 | 1639 | End point PCR |
| S1_PyR3 | AAATCGCAAGGCTAATGAATCC |  |  |  |
| Cloning analysis primer for SEGS-1 (NEB #E1202S) | |  |  |  |
| S1512A | ACCTGCCAACCAAAGCGAGAAC | 55 | >1639 | Colony PCR and  Sanger sequencing |
| S1513A | TCAGGGTTATTGTCTCATGAGCG |  |  |  |
| Overlapping sequencing primers for SEGS-1 | |  |  |  |
| S1-hp1F* | TACGCAGCAGCCATCATCGACATC |  |  |  |
| S1-2F * | GCAGTTCAGCAGTTCAACTG |  |  |  |
| S1-2R* | CAGTTGAACTGCTGAACTGC |  |  |  |
| SII 3F | AGGACCTTTGGAGCTCGA |  |  |  |
| S1-4F* | GGGTAGCCTCTAATCCTTCA |  |  |  |
| SII B5R | AAGCTTTACAAAGCTCAGCTTGGA |  |  |  |
| SII 6F | GATAACTTCCTAACTGGCTGC |  |  |  |
| S1-6R* | GCAGCCAGTTAGGAAGTTATC |  |  |  |
| S1_PyR4 | AAATTGGAACCACCACTCCA |  |  |  |
| Amplification and sequencing of CMD2-associated mutations^†^ | | | | |
| G680V-F | ACTATTAGCTGCTCGAAGAAGAG | 55 | 371 | G680V & A684G |
| G680V-R | ATCCACTTGATGGCGTATAACT |  |  |  |
| CMD2snEF | AATGCAGAAACTAGGAGGCGGCTT | 55 | 575 | V528L |
| CMD2snER | GGTTCTTCTGTTTAGCCTTCCT |  |  |  |
| V528L Seq2 | TGAAATGTAGTGAGTCTGCTACCTGT | |  | V528L sequencing |

*****Ndunguru et al., 2016

^†^Lim et al. 2022

**Table 3.** SEGS-1 fragments

| **Clone**  **name** | **SEGS-1**  **fragment** | **SEGS-1 coordinates** | | **Primer**  **name** | **Primer sequence** | **Annealing**  **(°C)** |
| --- | --- | --- | --- | --- | --- | --- |
|  |  | **Genomic** | **Clone** |  |  |  |
| pNSB | G | 1–277 | (+3)1–277 | SEG1FWDNot1 | GAATGCGGCCGCCACTACGCTACGCAGCAGCC | 63 |
|  |  |  |  | S1GnnRNot1 | GAATGCGGCCGCAGAGGAGCGAGGTGGGTG |  |
|  | H | 644–756 | 644-756 | S1InnFwdNot1 | GAATGCGGCCGCTTTACAAAGCTCAGCTTGG | 55 |
|  |  |  |  | SATII 1RevNot1 | GAATGCGGCCGCTGAAGGATTAGAGGCTACCC |  |
|  | J | 275–646 | 275–646 | SATII FwdNot1 | GAATGCGGCCGCGCAGTTCAGCAGTTCAACTG | 48 |
|  |  |  |  | SATII RNot1 | GAATGCGGCCGCGCTTTTATTAAAAAATTTTGG |  |
|  | F | 757–1007 | 757–1007(+13) | S1FnnFwdNot1 | GAATGCGGCCGCACTCTATTTTTCCGTTTGG | 52 |
|  |  |  |  | SEGS1REVNot1 | GAATGCGGCCGCGTACCACTACGCTACGCAG |  |

**SEGS-1 Fragments**

**G**

ACGCTACGCAG**|**CAGCCATCATCGACATCGTATTTTAA*CCAGAGGACCCGTCGACCGCCTGAGCAGCAGCACGTCGCACCAGCACCACCGCCGCATCGCGCGCCTGTGAGCCGCCGCACCACTGGATCTCGTGCTCGTGAGCCGCCGCACGCCGCAACTCTTCATCTACCGCTCGTTTACAGCCCACCTCTGTATCACGCGATTGTGAGCCGCCGACTGCCCGCCGCACGCCCGCACCTCTGCATCAACTGCTCGTTTGCCACCCACCTCGCTCCTCT*

**J**

*CTGC*AGTTCAGCAGTTCAACTGTAAGCATTTTTTCGTTAAATCTGAAGAAAATAGTTCTGGATAGAATTTTGATTGGTAAGCATTATGAATTTATTATGACATTCAAGTTTATAGGCATCATAGTGTTGCTTAGGACATACTTAGCTTGTAGTTCCAGAAAATAGAGTCATTTCTGGTTTTCTTTTACAATGGAGGTGTTTATTCCATTGTAATTTTGAGCTGAGCTTTGTTAAGGACCTTTGGAGCTCGAGCTTTGTTTACAAGGCATCTTGATAGAGCTTTTCGAGCTCGAATTAGAATTAGGCTCATGGTTATACTAAAGGGAGTTTTTCATGAGTTTGAGTGCTTCCAAAATTTTTTAATAAAAGC

**H**

GCTTTACAAAGCTCAGCTTGGATCGATTACACCTCTACTGACCCTACTCAGTTTGGGACTCTGGCTGGGGCCATTCTCAAAAGCCATTTATCTGGGTAGCCTCTAATCCTTCA

**F**

ACTCTATTTTTCCGTTTGGTTCTGAGAGAGTACTAAAAAGGAAATCCAACCATATATGATCAAATCTAATGATATAGCTGGTGAGTACTGCAACATAATTGCAATTTATGCAGTTATTTCTCTTGAATTTGGTATCTGCAATTTATGTATAAATCCCTAGCAGAATATTTTACTGGAGTGGTGAATATGTGTAGGCTTCACTATGGTGGAAATGGAAATTTGTGTGTGATAACTTCCTAACTGGCTGCT**|**GC

**N**

ACTCTATTTTTCCGTTTGGTTCTGAGAGAGTACTAAAAAGGAAATCCAACCATATATGATCAAATCTAATGATATAGCTGGTGAGTACTGCAACATAATTGCAATTTATGCAGTTATTTCTCTTGAATTTGGTATCTGCAATTTATGTATAAATCCCTAGCAGAATATTTTACTGGAGTGGTGAATATGTGTAGGCTTCACTATGGTGGAAATGGAAATTTGTGTGTGATAACTTCCTAACTGGCTGCT**|**CAGCCATCATCGACATCGTATTTTAA*CCAGAGGACCCGTCGACCGCCTGAGCAGCAGCACGTCGCACCAGCACCACCGCCGCATCGCGCGCCTGTGAGCCGCCGCACCACTGGATCTCGTGCTCGTGAGCCGCCGCACGCCGCAACTCTTCATCTACCGCTCGTTTACAGCCCACCTCTGTATCACGCGATTGTGAGCCGCCGACTGCCCGCCGCACGCCCGCACCTCTGCATCAACTGCTCGTTTGCCACCCACCTCGCTCCTCT*

**Fig. X** SEGS-1 fragments**.** The sequences of the SEGS-1 fragments that were assayed for activity are shown. The green shading indicates the GC-rich region. The blue line marks the junction position. The unlined nucleotides overlap the adjacent fragment.
